# Cigarette smoke induces pulmonary arterial dysfunction through an imbalance in the guanylyl cyclase redox status

**DOI:** 10.1101/2022.02.21.481310

**Authors:** J. Sevilla-Montero, J. Pino-Fadón, O. Munar-Rubert, M. Villegas-Esguevillas, B. Climent, M. Agrò, C. Choya-Foces, A. Martínez-Ruiz, E. Balsa, C. Muñoz, RM. Gómez-Punter, E. Vázquez-Espinosa, A. Cogolludo, MJ. Calzada

## Abstract

Chronic obstructive pulmonary disease (COPD), whose main risk factor is cigarette smoking (CS), is one of the most common diseases globally. Many COPD patients also develop pulmonary hypertension (PH), a severe complication that leads to premature death. Evidence suggests reactive oxygen species (ROS) involvement in COPD and PH, especially regarding pulmonary artery smooth muscle cells (PASMC) dysfunction. However, the effects of CS on the pulmonary vasculature are not completely understood. Herein we provide evidence on the effects of CS extract (CSE) exposure on PASMC regarding ROS production, antioxidant response and its consequences on vascular tone dysregulation. Our results indicate that CSE exposure promotes mitochondrial fission, mitochondrial membrane depolarization and increased mitochondrial superoxide levels. However, the increase in superoxide did not parallel a counterbalancing antioxidant response in human pulmonary artery (PA) cells. Interestingly, the mitochondrial superoxide chelator mitoTEMPO reduced mitochondrial fission and membrane potential depolarization caused by CSE. As we have previously shown, CSE reduces PA vasoconstriction and vasodilation. In this respect, mitoTEMPO prevented the impaired nitric oxide-mediated vasodilation, while vasoconstriction remained reduced. Finally, we observed a CSE-driven downregulation of the Cyb5R3 enzyme, which prevents soluble guanylyl cyclase oxidation in PASMC. This might explain the CSE-mediated decrease in PA vasodilation. These results provide evidence that there might be a connection between mitochondrial ROS and altered vasodilation responses in PH secondary to COPD, and strongly support the potential of antioxidant strategies specifically targeting mitochondria as a new therapy for these diseases.

**Graphical abstract:** 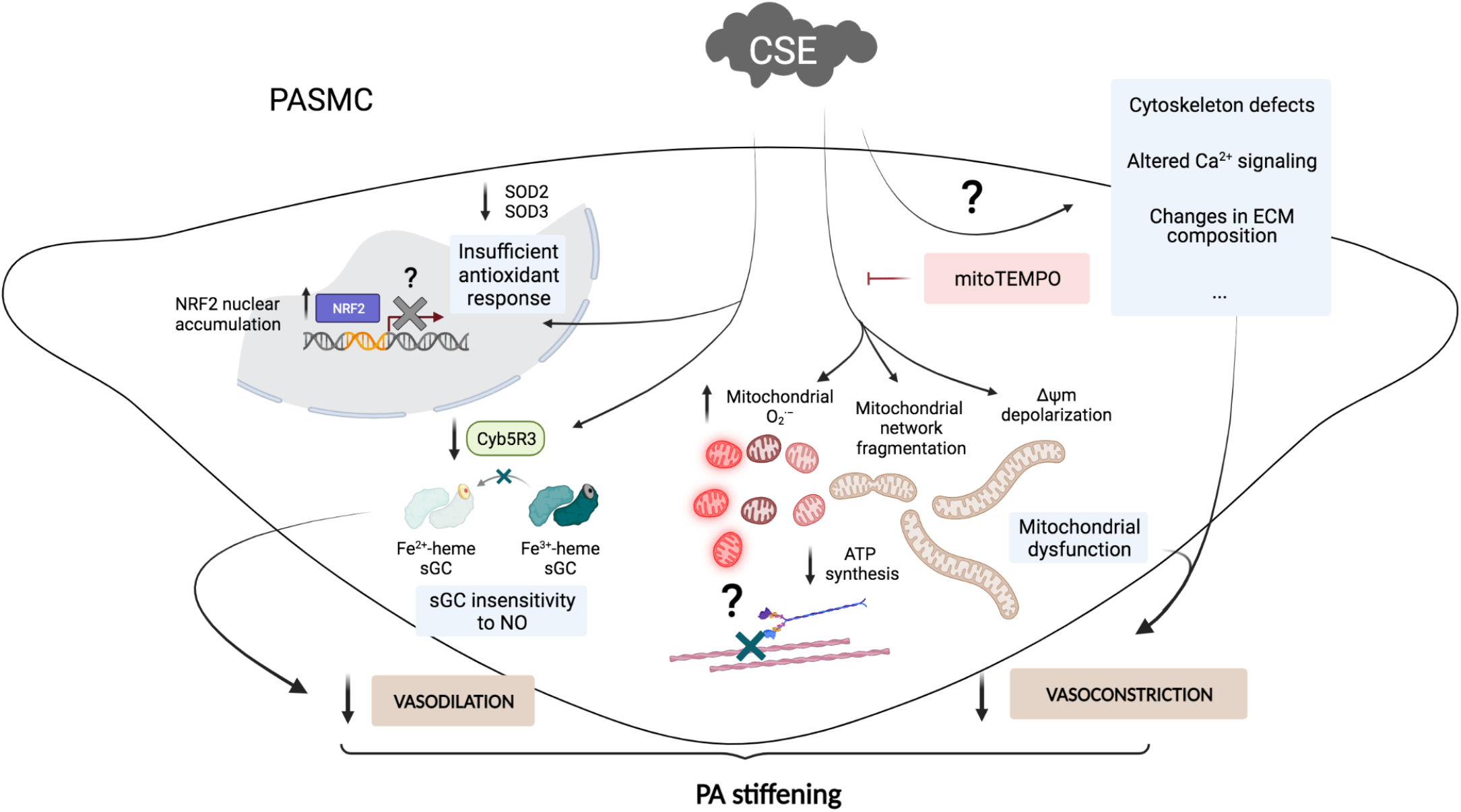

## INTRODUCTION

Chronic obstructive pulmonary disease (COPD), a pathology characterized by chronic bronchitis, emphysema, small airway obstruction and persistent respiratory symptoms, is strongly influenced by environmental agents, primarily cigarette smoking (CS)^1^. In addition, some COPD patients suffer pulmonary hypertension (PH) due to the vascular remodeling of their pulmonary arteries (PA), with characteristic vascular wall hypertrophy and vascular tone dysregulation, resulting in increased pulmonary artery pressure (PAP) and right ventricle alterations^2,3^. In spite of being such alarming diseases, curative therapies for COPD or PH are not available yet and a more comprehensive understanding of their underlying mechanisms is needed to develop new therapeutic approaches.

Although the CS-driven damage on the airways has been extensively studied, its direct effects on the pulmonary vasculature are not so well known, mainly because vascular remodeling was thought to just be a consequence of alveolar hypoxia due to emphysema and airway obstruction. However, in patients with severe emphysema and airway obstruction, PAP only slightly correlates with hypoxemia and airflow limitation, and no correlation exists between PAP and emphysema severity^4,5^. Furthermore, media thickening is also observed in PA of mild COPD patients and smokers without airway obstruction^6^. This suggests that other unknown mechanisms, hypoxia-independent and directly triggered by CS on the pulmonary vasculature, may also be responsible for PH development on these patients^7^. In fact, our recently published results demonstrate that cigarette smoke extract (CSE) causes a decrease of PA vasoconstriction and vasodilation responses, with alterations intrinsic to the smooth muscle cell (SMC) layer which could contribute to CS-dependent PH development^8^.

COPD development is closely related to oxidative stress within the lung. This has been identified in terms of augmented oxidative stress markers in exhaled breath condensates of COPD patients compared to control subjects^9–14^, but also at the lung tissue level^15–21^. It has been stablished that a main source of reactive oxygen (ROS) in the lung arises from CS and air pollutants themselves, especially unsaturated aldehydes such as acrolein^22^. However, endogenously-produced oxidant molecules also play an important role in COPD pathogenesis, where systems like NADPH oxidase (NOX), xanthine oxidase and mitochondrial respiration appear to be the most relevant sources of endogenous ROS^23^. Conversely, antioxidant defenses unable to counteract the burden in oxidant molecules accumulation have also been linked to COPD and its exacerbations, like reduced availability of glutathione^24^. Additional evidence include the downregulation of nuclear erythroid-2 related factor 2 (NRF2), the main transcription factor promoting endogenous antioxidant responses^25,26^, the depletion of other antioxidants such as vitamins A and E, and the alterations of DNA repair mechanisms^27–29^.

Among the different endogenous sources of pulmonary ROS, NOX family comprises NOX1-5 transmembrane enzymes, which catalyze electron transfer from NADPH+H^+^ to molecular oxygen to form superoxide and/or hydrogen peroxide^30^. These core subunits need to associate to transmembrane protein p22^phox^, as well as to a cytosolic group of cofactors that translocate to the membrane to stimulate ROS production. NOX2-related cofactors are p47^phox^, p40^phox^, p67^phox^, whereas cytosolic subunits associated to NOX1 are NOX organizer 1 (NOXO1) and NOX activator 1 (NOXA1). NOX4 is also p22^phox^-dependent but does not seem to require any cytosolic subunits, so it is thought to be constitutively active^30^.

Higher amounts of NOX2-expressing immune cells are found within the lung of COPD patients, contributing to the inflammation and oxidative stress of the surrounding tissue^31^, but NOX4 levels in airway epithelial and SMC also correlate with COPD severity^32–34^. Interestingly, Guo *et al* also found that pulmonary artery smooth muscle cells (PASMC) from COPD patients have higher NOX4 levels^35^, although if this condition arises from CS direct impact or hypoxia contribution is unclear. Evidence suggests that CS might directly take a part on NOX-derived vascular oxidative stress, since rat aortic SMC show higher NOX1 levels responsible for increased superoxide upon *in vitro* CSE exposure^36^.

Dysfunctional mitochondria are also important superoxide sources, as a side product of their respiratory chain activity, mainly through ROS release towards the mitochondrial matrix or to the intermembrane space^37^. It is commonly accepted that increased ROS production in the mitochondria is mainly associated to higher values of mitochondrial membrane potential (Δψ_m_) and a slow electron transference along the respiratory chain. Mild uncoupling thus helps to reduce this superoxide production^38^, although there exist situations where this uncoupling does not reduce mitochondrial ROS, in spite of decreasing Δψ_m_^39,40^. Redox-optimized ROS balance hypothesis was thus developed to reconcile this paradox and take into account both mitochondrial ROS production and the redox status of molecules involved in antioxidant mechanisms, such as NADPH+H+ or glutathione^41,42^. Mitochondrial damage induced by CS exposure in COPD leads to the accumulation of dysfunctional mitochondria with decreased ATP synthesis and increased ROS production rates ^43^, which may also play a key role in the vascular remodeling during COPD pathogenesis.

Cellular ROS levels are also delimited by a group of antioxidant enzymes regulated by NRF2, a transcription factor encoded by *NFE2L2* gene^44^. Interestingly, a study on the association between NRF2 gene polymorphisms and airflow insufficiency in smokers points out that defective NRF2 might play a role in COPD due to excessive oxidative burden and insufficient antioxidant, protective responses^45^. Additionally, decreased *NFE2L2* mRNA and NRF2 protein levels are detected in lung tissue, pulmonary macrophages and bronchial epithelium of COPD patients compared to healthy donors^26,46–48^. Despite this, CS role in NRF2 pathway disruption is unclear, because several groups have shown that, at least *in vitro*, CSE drives NRF2 total upregulation and nuclear translocation in human bronchial epithelial cells^49–51^. Therefore there is no clear evidence on the role of NRF2 in COPD-related PH.

ROS can also react with nitric oxide (NO), generating on the one hand reactive nitrogen species such as peroxynitrite^52^, and on the other hand scavenging NO levels and diminishing its downstream effects in the vasculature. NO produced by endothelial NO synthase (eNOS) diffuses to reach SMC, where it activates soluble guanylate cyclase (sGC) to synthesize cGMP and stimulate protein kinase G to promote vasodilation. CSE effects on pulmonary NO pathway are mostly characterized in terms decreased NO production^53–55^. Accordingly, endothelium-dependent PA vasodilation is reduced in different animal models upon CS challenge^56–58^. However, direct effects of CS on PA tunica media regarding this pathway are not fully understood, and evidence suggesting additional molecular dysfunction of sGC or other downstream NO effectors in SMC are still unclear.

sGC enzyme comprises a heterodimer of α and β subunits, where sGCβ subunit binds a heme cofactor containing a Fe^2+^ ion which coordinates to NO^59^. Fe^2+^ ion is fundamental for NO binding, since its oxidized form is no longer sensitive to NO^60^. Cytochrome b5 reductase 3 (Cyb5R3) has been identified as responsible for maintaining this reduced Fe^2+^ status of sGC in vascular SMC^61^, and mice lacking SMC *Cyb5r3* expression show higher arterial pressure values and are more sensitive to angiotensin-driven hypertension^62^.

Compounds with the ability of modifying sGC activity are grouped into two main families with different pharmacological properties: sGC stimulators (such as riociguat, with upregulates sGC activity when the heme group is in its reduced, Fe^2+^ state) and sGC activators (like ataciguat or cinaciguat, which promote sGC activity on its oxidized and even heme-free forms)^63^. These compounds have shown their benefits in COPD-related PH, since sGC stimulators riociguat and BAY 41-2272 prevent right ventricle hypertrophy and vascular muscularization in mouse and guinea pig models of chronic CS exposure^64^. Nonetheless, since these compounds also prevent emphysema development and induce bronchodilation^64,65^, NO-sGC pathway dysfunction still cannot be unequivocally assigned to hypoxia, CS or a combination of both.

In the present study, we analyzed the CSE *in vitro* effects related to oxidative stress accumulation and NO-sGC vasodilation pathway in human PA smooth muscle cells (hPASMC). Particularly, we observed a CSE-mediated increase in mitochondrial superoxide levels, which was not accompanied by an upregulation of antioxidant enzymes. This mitochondrial ROS burden within PASMC likely targets NO-sGC signaling pathway, which explains the prevention of CSE-mediated dysfunction in the presence of mitoTEMPO. Additionally, our results pointed to a significant downregulation of Cyb5R3, which might promote an excessive oxidation of the sGC, as suggested by PA vasodilation being diminished in response to riociguat and conversely augmented in response to ataciguat.

## MATERIALS AND METHODS

### Cell culture

Primary human pulmonary artery smooth muscle cells (hPASMC) were obtained from ScienCell Research Laboratories (#3110). Cells were cultured, following manufacturer’s recommended specifications, in smooth muscle cell complete medium (ScienCell, #1101) containing 2% fetal bovine serum (FBS, ScienCell, #0010), 1X smooth muscle cell growth supplement (ScienCell, #2352 and #1152), 100 U/mL penicillin and 100 μg/mL streptomycin (ScienCell, #0503). Cells were used for a maximal number of 8 passages and were maintained at 37 °C in a humidified atmosphere of 5% CO_2_. For some experiments, cells were treated with 25 nM mitoTEMPO (Sigma, SML0737).

### Cigarette smoke extract (CSE) preparation

CSE was prepared from commercial Marlboro Red cigarettes (Philip Morris Brand Sàrl Neuchâtel, Switzerland, 13 mg tar and 1.0 mg nicotine/cigarette^66^). The smoke from the complete consumption of five cigarettes was continuously bubbled into 150 mL of PBS using VACUSAFE aspiration system (INTEGRA Biosciences AG) at a vacuum pump flow rate of 8 L/min. CSE solution was filtered through a 0.20 μm-pore system, immediately aliquoted and kept at −80 °C until used. To ensure a similar preparation amongst different batches, CSE concentration was measured spectrophotometrically at 320 nm wavelength. This solution was considered to be 100% CSE and diluted to obtain the desired concentrations for each experiment. To avoid the exposure to volatile substances from the smoke, untreated and CSE-challenged cells were always kept in different incubators.

### Mitochondrial mass and mitochondrial superoxide levels quantification

Mitochondrial mass levels and specific mitochondrial superoxide levels in the absence or presence of mitoTEMPO were quantified with the fluorescent probes MitoTracker™ Green FM (ThermoFisher Scientific, M7514) and MitoSOX™ Red (ThermoFisher Scientific, M36008). Following 24-hour treatment with CSE, cells were incubated with DMSO as vehicle control, or with 25 nM MitoTracker™ Green FM and 5 μM MitoSOX™ Red, in HBSS for 15 minutes at 37 °C. Afterwards, cells were trypsinized and median MitoTracker™ Green FM and MitoSOX™ Red fluorescence was measured on a FACSCanto™ II cytometer (BD Biosciences) illuminating with Ar 488 nm laser and excluding cell debris and doublets from the analysis. Specific changes in MitoTracker™ Green FM and MitoSOX™ Red signal intensities were quantified by subtracting fluorescence from DMSO-treated hPASMC, and normalized to non CSE-exposed cells.

### Western blot analysis

Cells were grown to 90% confluence in 6-well plates in the presence or absence of CSE for 24 hours and lysates were prepared in non-reducing 2X Laemmli buffer, boiled at 95 °C for 10 minutes in the presence of 20 mM DTT, electrophoretically separated by SDS-PAGE and transferred onto 0.45-μm nitrocellulose membranes (GE Healthcare Life Sciences, 10600003). Total protein bands were reversibly stained with Fast Green FCF (Sigma, F7252) and imaged on ImageQuant LAS 4000 or Amersham ImageQuant 800 (GE Healthcare Life Sciences) for total protein quantification. Transferred proteins were probed overnight at 4 °C with specific primary antibodies (Table 1). Horseradish peroxidase-conjugated secondary antibodies (anti-mouse IgG, Dako, P0260, and anti-rabbit IgG, Invitrogen, 32460) were added for 1 hour at room temperature and protein signal was then visualized using Immobilon Forte (Millipore, WBLUF0500) on ImageQuant LAS 4000 or Amersham ImageQuant 800. Specific protein bands intensity was quantified by densitometry using ImageJ 1.51 software (U. S. National Institutes of Health, Bethesda, Maryland, USA) and normalized to the intensity of Fast Green FCF staining from each complete gel lane.

**Table 1.** List of antibodies used for western blot and their corresponding product information.

| Target protein | Clonality (clone) | Host species | Manufacturer (catalog number) |
| --- | --- | --- | --- |
| SOD2 | Monoclonal (mAbcam74231) | Mouse | Abcam (ab74231) |
| SOD1 | Monoclonal (24) | Mouse | Santa Cruz (sc-101523) |
| SOD3 | Monoclonal (4G11G6) | Mouse | Abcam (ab80946) |
| catalase | Monoclonal (A-4) | Mouse | Santa Cruz (sc-271358) |
| NRF2 | Monoclonal (383727) | Mouse | R&D Systems (MAB3925-SP) |
| α-tubulin | Monoclonal (DM1A) | Mouse | Sigma (T9026) |
| H2A | Monoclonal (L88A6) | Mouse | Cell Signaling Technology (3636) |
| NOX1 | Polyclonal | Rabbit | Novus (NBP1-31546) |
| NOX4 | Polyclonal | Rabbit | Novus (NB110-58849SS) |
| p22phox | Polyclonal | Rabbit | Abcam (ab75941) |
| NOXA1 | Monoclonal (H-6) | Mouse | Santa Cruz (sc-398873) |
| NOXO1 | Monoclonal (F-5) | Mouse | Santa Cruz (sc-390927) |
| sGCα | Polyclonal | Rabbit | Abcam (ab50358) |
| sGCβ | Polyclonal | Rabbit | Sigma (SAB4501344) |
| Cyb5R3 | Polyclonal | Rabbit | ProteinTech (10894-1-AP) |

### Cytosolic and nuclear subcellular fractionation

For cytosolic and nuclear protein fractions isolation, NE-PER™ Extraction Reagent (Pierce, 78833) was used following manufacturer recommendations. Cells were grown to 90% confluence in 100 mm plates in the presence or absence of CSE for 24 hours (two 100 mm plates per condition). After CSE exposure, cells were trypsinized and split for both total protein lysate analysis and cytosolic/nuclear extraction using NE-PER™ kit. Total, cytosolic and nuclear fractions were solubilized in RIPA buffer and protein content was analyzed by bicinchoninic acid assay (ThermoScientific, 23225), prior to their analysis by western blot. Cytosolic and nuclear fractions purity was assessed by a-tubulin and H2A histone levels determination, respectively.

### Mitochondrial oxygen consumption

Oxygen consumption rate (OCR, pmol/min) of hPASMC were measured using Seahorse XF96 metabolic analyzer and Seahorse XF Cell Mito Stress Test kit (103015-100, Seahorse XF Technology, Agilent), following manufacturer’s recommendations. Cells (3 · 10^4^ hPASMC per well) were seeded onto XF96 V3 PS Cell Culture Microplates (101085-004, Agilent) and exposed to air or CSE for 24 hours before being loaded into the analyzer. OCR was monitorized over time while 5 μM oligomycin, 0.5 μM carbonyl cyanide-*p*-trifluoromethoxyphenylhydrazone (FCCP) and 2 μM rotenone/5 μM antimycin A from Mito Stress Test kit were sequentially added to the cells. Mitochondrial respiration-related parameters where quantified in terms of OCR as follows: basal respiration (OCR prior to oligomycin addition minus OCR after rotenone/antimycin A addition), H^+^ leakage (OCR prior to oligomycin addition minus OCR after oligomycin addition), maximal respiratory capacity (OCR after FCCP addition minus OCR after oligomycin addition), ATP-linked respiration (basal respiration minus H^+^ leakage OCR) and spare capacity (maximal capacity minus basal respiration).

### Mitochondrial network image analysis

Mitochondrial network morphology was assessed transfecting hPASMC grown on a 60 mm plate to 50% confluence with 1 μg pDsRed2-mito expression vector (Clontech Laboratories, 632421) and 2 μL Lipofectamine™ 2000 (ThermoFisher Scientific, 11668027). 24 hours after transfection, hPASMC were trypsinized, allowed to attach on 10 μg/mL fibronectin-functionalized 13-mm glass coverslips and challenged with CSE in the presence or absence of mitoTEMPO for 24 hours. Following CSE treatment, cells were fixed with 4% paraformaldehyde (Electron Microscopy Sciences, 15710) in PBS for 10 minutes at room temperature and mounted on slides in Faramount Aqueous Medium (Dako, S3025). Cells were imaged with a Leica DMR microscope using an immersion 100x objective (HCX PL APO 100X/1.40-0.7 oil CS, Leica) and illuminated with a mercury lamp (ebq 100, Lightning & Electronics JENA). Ten images per sample were collected using Leica DCF360FX camera and LAS V4.1 software, and mitochondrial network was analyzed following the previously described computational strategy^67^, using ImageJ 1.51 software. Images were pre-processed by means of unsharp masking (radius = 3 pixels), local contrast enhancing and median filtering (radius = 2 pixels) and, afterwards, a binary mask of every image was generated to measure the area covered by mitochondrial signal (mitochondrial footprint) and to generate a skeletonized representation of the network. This skeletonized image was characterized in terms of the number of individual mitochondria, number of networks (containing at least one junction pixel), mean length of network branches and mean number of branches per network. Number of individual mitochondria and networks parameters were normalized over the mitochondrial footprint from the same image.

### Δψ_m_ assessment

Following 24-hour exposure to CSE, Δψ_m_ in hPASMC was measured using the fluorescent probe tetramethylrhodamine methyl ester (TMRM, ThermoFisher Scientific, T7668), following the staining method described by Creed and McKenzie^68^. Cells were seeded on 8-well chambered coverslips (ThermoScientific, 155411) and stained with 20 nM TMRM and 2 μg/mL Hoechst in recording solution (RS, 109 mM NaCl, 50 mM KCl, 2 mM MgSO_4_, 1.25 mM KH_2_PO_4_, 10 mM D-glucose, 2 mM CaCl_2_ and 10 mM Hepes) for 40 minutes at 37 °C. Afterwards, cells were imaged on a Leica SP5 confocal microscope in the presence of 20 nM TMRM in RS using an immersion 40x objective (HCX PL APO CS 40X/1.25 oil UV, Leica) and illuminating with 488 nm Ar and 561 nm diode-pumped solid-state lasers. Cells were continuously recorded for 5 minutes before adding 10 μM FCCP (Sigma, C2920) to induce complete Δψ_m_ depolarization for an additional 5-minute lapse. Confocal images were collected using Leica TCS SP5 software (Leica Microsystems, Mannheim, Germany). Subsequent analysis was performed on ImageJ 1.51, subtracting TMRM-channel background by means of rolling ball filtering method (radius = 50 pixels) and quantifying mean TMRM fluorescence intensity of every cell along time.

### ATP levels quantification

Total ATP levels were quantified using Luminescent ATP Detection Assay Kit (ab113849, abcam) following manufacturer’s recommendations. After hPASMC challenge to CSE in the absence or presence of mitoTEMPO, cells were lysed for 5 minute and luciferin-luciferase solution was added during an additional 5-minute time lapse. Following this incubation, luminescence was registered on a Glomax microplate reader (Promega) and ATP concentration was extrapolated from a reference curve of known ATP standard dilutions. Results were normalized by total protein determination using bicinchoninic acid assay (ThermoScientific, 23225).

### Vascular contractility measurement

PA from WT C57BL/6J mice were carefully dissected free of surrounding tissue, cut into rings (1.8-2 mm length) and overnight-exposed to 10% CSE in the presence or absence of mitoTEMPO. Afterwards, vessel segments were mounted on a wire myograph in the presence of Krebs physiological solution. Contractility was recorded by an isometric force transducer and a displacement device coupled with a digitalization and data acquisition system (PowerLab, Paris, France). Preparations were firstly stimulated by rising the K^+^ concentration of the buffer to 80 mM in exchange for Na^+^. Vessels were washed three times and allowed to recover before a new stimulation. Arteries were then stimulated with increasing concentrations of serotonin (5-HT, from 10 ^-8^ to 3· 10 ^-5^ M, Sigma), endothelium-dependent vasodilator acetylcholine (ACh, from 10 ^-9^ to 10^-5^ M, Sigma) and NO donor sodium nitroprusside (SNP, from 10 ^-11^ to 10 ^-5^ M, Sigma). To evaluate sGC redox status, following PA vasoconstriction induction with 10^-5^ M 5-HT, vasodilation responses to increasing concentrations of the sGC stimulator riociguat or sGC activator ataciguat (both from 10^-9^ to 10^-5^ M) were tested.

### Statistical analysis

All data are presented as the mean ± standard error of the mean (SEM). For all of the analyses, normality condition was evaluated using Saphiro-Wilk test and, according to its results, subsequent analysis of the data was carried out as follows. On the one hand, if normality condition could be assumed, parametric statistical tests were used. When comparing the mean of a group and a reference value, two-tailed one-sample Student’s *t* test was used. Alternatively, when comparing the means of three or more groups, one or two-way analysis of variance (ANOVA) tests followed by Bonferroni’s pairwise comparisons or linear trend *post hoc* tests were used. Finally, repeated measures ANOVA test followed by Bonferroni’s *post hoc* pairwise comparisons was used to compare the means of three or more non-independent groups. On the other hand, if normality condition was not accomplished, non-parametric statistical tests were used, and Kruskal-Wallis’s *H* test followed by Dunn’s *post hoc* pairwise correction was used when comparing the medians of three or more groups. A p-value *P*<0.05 was considered statistically significant. All of the statistical analyses were performed on GraphPad Prism 7.0a software (San Diego, CA, USA).

## RESULTS

### CSE exposure results in increased mitochondrial superoxide levels and reduced antioxidant defenses in PASMC

Recent results from our laboratory indicate that CSE increases intracellular superoxide levels in human PA fibroblasts and hPASMC^8^, although the source of ROS in these cells is still unknown. Since one of the main vascular superoxide sources described in the lung and the pulmonary vasculature are the NOX complexes, we studied the expression of NOX1 and NOX4 enzymes and other additional subunits that contribute to their activity (namely p22^phox^, NOXO1 and NOXA1) in hPASMC challenged to CSE. Surprisingly, none of these proteins proved to be increased. Furthermore, following CSE-exposure we observed a statistically significant downregulation of p22^phox^ protein, the subunit anchoring both functional NOX1 and NOX4-containing complexes to the cell membranes (Figure 1A).

**Figure 1:**
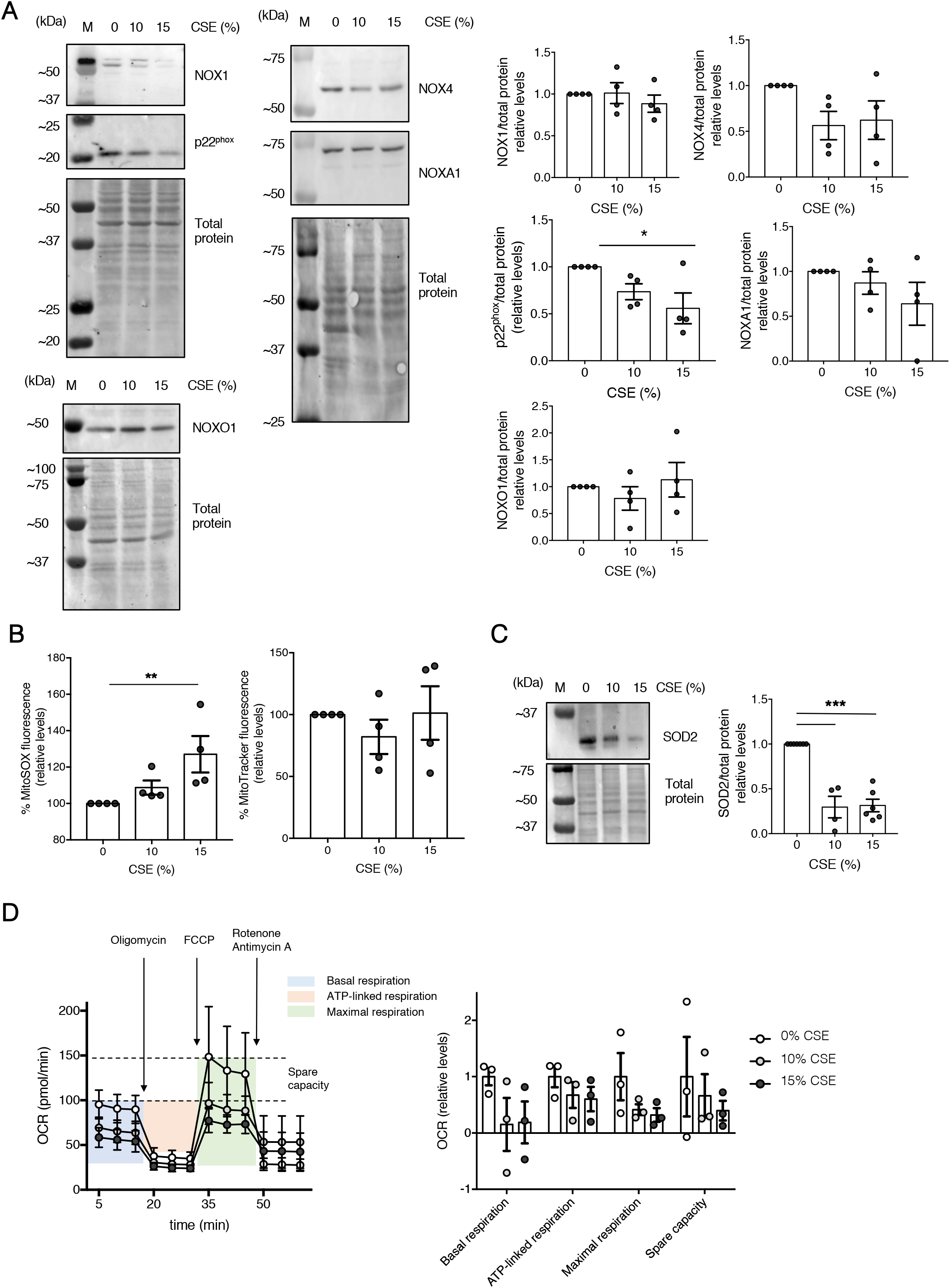
Impact of CSE on NOX subunits, mitochondrial superoxide, SOD2 enzyme levels and mitochondrial respiration functions. Following 24-hour challenge with CSE at the indicated concentrations, NOX subunits and SOD2 levels in hPASMC were analyzed by western blot, mitochondrial superoxide levels and mitochondrial mass were analyzed by flow cytometry, and mitochondrial respiration was analyzed in terms of oxygen consumption rates (OCR). **A.** Protein levels of hPASMC were analyzed by western blot probed against NOX1, p22^phox^, NOX4, NOXA1, NOXO1 and total protein. Representative images (left) and band quantifications by densitometry (right) are shown. M: molecular weight marker lane. Densitometry data were controlled with total protein staining as loading control, expressed as fold change over 0% CSE-exposed cells, and presented as mean ± SEM; n = 4. Statistical comparisons between groups were made using one-way ANOVA test followed by Bonferroni’s *post hoc* test (\**P*<0.05). **B.** Mitochondrial superoxide (left) and mitochondrial mass (right) levels were quantified using 5 μM MitoSOX™ and 25 nM MitoTracker™ probes, respectively. Values were calculated as the median fluorescence intensity of MitoSOX™ or MitoTracker™-stained cells minus the median fluorescence intensity of unstained cells, expressed as percentages over 0% CSE-exposed cells, and presented as mean ± SEM; n = 4. Statistical comparisons among groups were made using Kruskal-Wallis’ *H* test followed by Dunn’s *post hoc* test (\*\**P*<0.01). **C.** Protein levels of hPASMC were analyzed by western blot and probed against SOD2 and total protein. A representative image (left) and band quantifications by densitometry (right) are shown. M: molecular weight marker lane. Densitometry data were controlled with total protein staining as loading control, expressed as fold change over 0% CSE-exposed cells, and presented as mean ± SEM; n = 4-6. Statistical comparisons among groups were made using oneway ANOVA test followed by Bonferroni’s *post hoc* test (\*\*\**P*<0.005). **D.** Mitochondrial respiration functions of hPASMC were analyzed quantifying OCR values along time (left), mean ± SEM; n = 3. Basal respiration, ATP production, maximal respiration and spare capacity values (right) calculated from OCR measurements, expressed as fold change over 0% CSE-exposed cells and presented as mean ± SEM; n = 3. Statistical comparisons among groups were made using one-way ANOVA test followed by Bonferroni’s *post hoc* test.

Since NOX enzymes did not seem to be specially relevant in this context, we then tried to characterize instead the mitochondrial contribution to oxidative stress in hPASMC challenged with CSE, and quantified mitochondrial superoxide levels by flow cytometry. We observed a significant increase in mitochondrial superoxide levels that were not accompanied by an increment in total mitochondrial mass, as measured by MitoSOX and MitoTracker staining respectively (Figure 1B). Most importantly, our results proved a significant CSE-mediated downregulation on mitochondrial superoxide dismutase 2 (SOD2) protein levels in hPASMC (Figure 1C), which might further contribute to increase mitochondrial superoxide levels.

Next, we studied the effects of CSE in the electron transport chain, as it is considered a major source of mitochondrial ROS. We then quantified hPASMC mitochondrial respiration rate by means of Seahorse assays to address if this mitochondrial superoxide accumulation would actually be associated to an organelle dysfunction. Our data suggested that CSE induced a decrease of hPASMC mitochondrial respiration following CSE exposure, as quantified in terms of a trend towards the reduction in the oxygen consumption rates associated to basal respiration, ATP-linked respiration, maximal respiration and spare capacity when compared to non-CSE exposed hPASMC (Figure 1D).

Oxidative stress might also arise from an unbalanced equilibrium between pro-oxidant and antioxidant elements, we tested if CSE challenge also compromised global antioxidant responses besides SOD2 downregulation. Our results proved that while no changes in cytosolic SOD1 levels were found, CSE significantly diminished extracellular SOD3 and catalase enzymes (Supplementary figure 1A). To further characterize this anomalous cell behavior against a supposedly pro-oxidant stimulus like CSE, we considered evaluating the levels of NRF2, which accumulates in the nucleus upon oxidative stress to promote antioxidant gene expression^69^. Following hPASMC 15% CSE exposure, we assessed NRF2 protein levels, in both cytosolic and nuclear fractions by western blot. Our results showed, however, that NRF2 nuclear accumulation could indeed be taking place in CSE-challenged hPASMC compared to control, unexposed ones (Supplementary figure 1B).

### Mitochondrial superoxide scavenger mitoTEMPO reduces CSE-driven mitochondrial fission in hPASMC

Our data pointed towards a dysfunctional oxidative phosphorylation being included among the different effects exerted by CSE on hPASMC, resulting in the overproduction of superoxide free radical. Thus, this superoxide might contribute to the phenotype of aberrant vasoactive responses of the PA that we have previously observed following CS exposure^8^. Different mitochondria-targeted antioxidants, such as mitoQ or mitoTEMPO, have shown beneficial effects on reducing mitochondrial superoxide levels on different models of pulmonary damage^70–72^. To evaluate if such an approach could also be valid in the pulmonary vascular cells context, hPASMC were incubated with CSE in the presence or absence of mitochondrial superoxide chelator mitoTEMPO. As expected, treatment with mitoTEMPO significantly attenuated the CSE-mediated increase on superoxide in these cells (Supplementary figure 2A).

Mitochondrial fission and fusion dynamics are fundamental cell strategies to continuously self-renew dysfunctional mitochondria upon exogenous stress insults. CSE is known to induce fragmentation of the mitochondrial network within airway SMC and epithelial cells resulting from an abnormal balance between fusion and fission mechanisms^73–75^. To analyze the morphological effects that CSE had on hPASMC mitochondrial network, and whether or not they could be reverted with mitoTEMPO treatment, we first transiently transfected the cells with a vector encoding for mitochondria-targeted DsRed2 protein. CSE-treated cells expressing this fluorescent protein showed a distinctive distribution throughout their mitochondrial networks (Figure 2A). This allowed us to image and transform this networks onto binary, skeletonized shapes (Supplementary figure 2B-C), and describe them in terms of individualized or network-forming mitochondria, as well as quantify the percentage of image covered by DsRed2 signal (namely, mitochondrial footprint), the average length of network-forming branches and, finally, the average number of branches per mitochondrial network. Using this image analysis tool, we compared the effects of CSE and mitoTEMPO onto hPASMC mitochondria. Data showed that CSE induced mitochondrial network fragmentation (Figure 2A, left), resulting in a significant increase in the number of individualized mitochondria (Figure 2A, left, white arrows), together with a slight increase in the number of mitochondrial networks as well (Figure 2A, right). Although average length of the different branches did not significantly change upon CSE-challenge, networks in general were comprised of a lesser number of branches compared to non-CSE-exposed cells (Figure 2A, right). Globally, all four parameters were compatible with a situation where, with no apparent changes in total mitochondrial mass, this was being restructured and preferentially relocated on individualized mitochondria rather than connected, fused networks. Interestingly, mitoTEMPO seemed to partly revert some of these CSE-driven effects on the mitochondrial network morphology, especially on the amount of individualized and network-forming mitochondria (Figure 2A, right).

**Figure 2:**
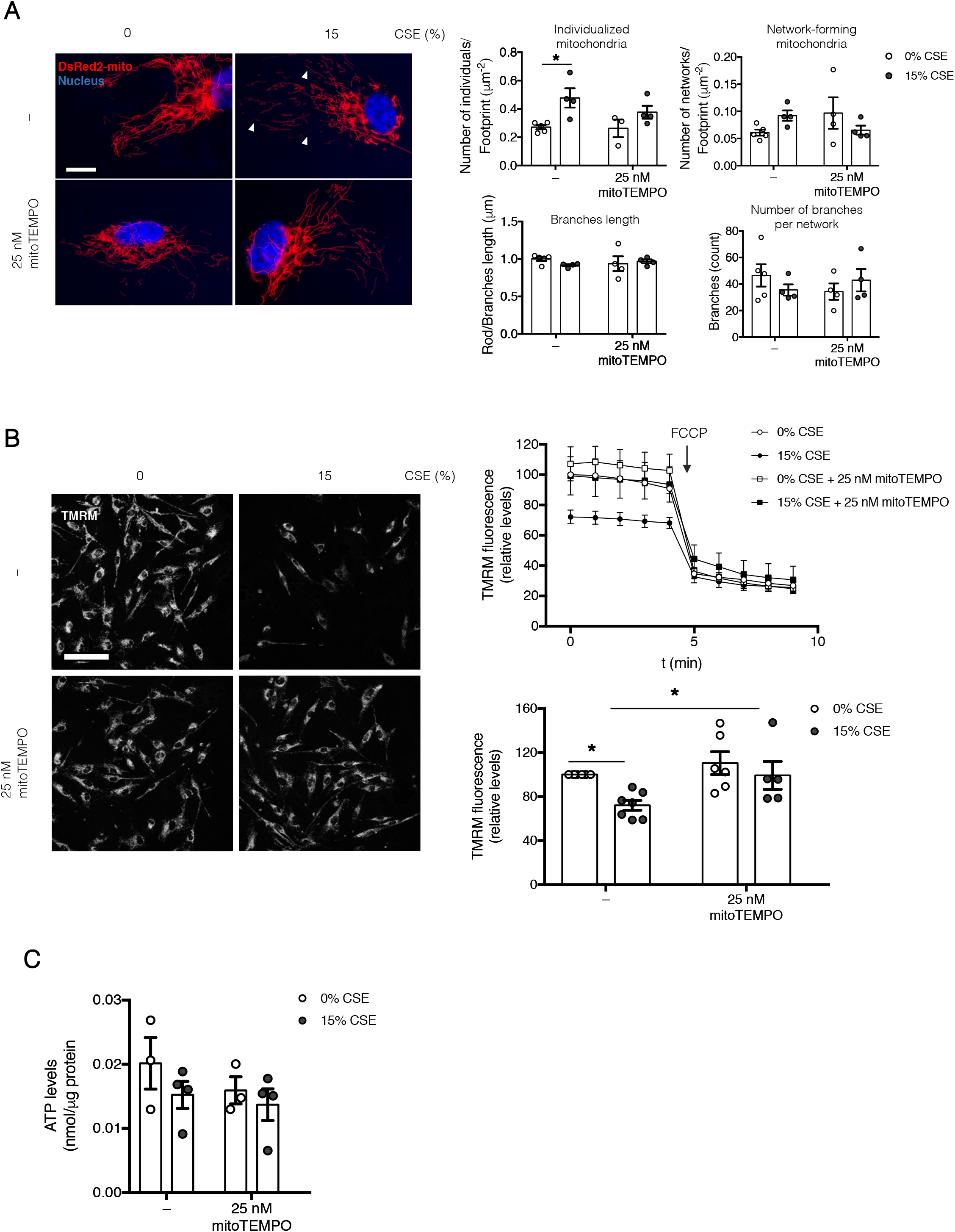
Effects of mitoTEMPO treatment on CSE-triggered mitochondrial network fragmentation and Δψ_m_ depolarization in hPASMC. Following 24-hour challenge with CSE in the absence or presence of 25 nM mitoTEMPO, hPASMC mitochondrial network morphology, mitochondrial membrane potential (Δψ_m_) values and total ATP levels were analyzed. **A.** Left: representative images of pDsRed2-mito transfected hPASMC, showing merged DsRed2-mito (red) and Hoechst (blue) signals, scale bar = 10 μm. White arrowheads indicate fragmented mitochondrial networks. Right: individualized and network-forming mitochondria values normalized over the surface covered by pDsRed2-mito signal (mitochondrial footprint) are shown, together with mean branches length within networks and average number of branches per network quantification. All values are presented as mean ± SEM; n = 3-5. Statistical comparisons among groups were made using two-way ANOVA test followed by Bonferroni’s *post hoc* test (\**P*<0.05). **B.** Left: Confocal lifeimaging was used to quantify Δψ_m_ in hPASMC stained with 20 nM TMRM; representative images are shown, scale bar = 100 μm. Right: TMRM fluorescence intensity values were calculated for 5 min before and 5 min after 10 μM FCCP addition, as Δψ_m_ depolarization positive control, from a minimal number of 5 cells per condition. Values were expressed as percentages over the first time point of 0% CSE-exposed cells in the absence of mitoTEMPO, and presented as mean ± SEM; n = 5-7 (top). Statistical comparisons of only the first TMRM intensity value among groups were made using two-way ANOVA test followed by Bonferroni’s *post hoc* test (\**P*<0.05) (bottom). **C.** Total ATP levels were quantified by chemiluminescence and normalized over total protein amount quantified by bicinchoninic acid assay. Values are presented as mean ± SEM; n = 3-4. Statistical comparisons were made using two-way ANOVA test followed by Bonferroni’s *post hoc* test.

### MitoTEMPO protects hPASMC from mitochondrial membrane depolarization caused by CSE

Beyond these morphological criteria, we aimed to further analyze the protective effects of mitoTEMPO, regarding a characterization of hPASMC mitochondria from a functional point of view. Given our observations of reduced mitochondrial respiration rate upon CSE treatment, this would likely translate into abnormal Δψ_m_ values and associate with reduced ATP levels. Therefore, after CSE exposure in the absence or presence of mitoTEMPO, we quantified Δψ_m_ status using life-imaging of TMRM-stained hPASMC (Figure 2B). As expected, CSE significantly induced a mitochondrial depolarization on hPASMC, as shown by reduced TMRM fluorescence compared to non-exposed cells (Figure 2B). Interestingly, and congruent with a less fragmented, healthier mitochondrial network, mitoTEMPO induced a significant recovery of this decreased Δψ_m_ due to CSE exposure (Figure 2B).

Finally, we wondered if this change in Δψ_m_ induced in hPASMC challenged to CSE affected the ATP cellular pool. Although our results did not reach statistical significance, they point towards a slight decrease in ATP levels upon acute CSE exposure (Figure 2C), which is in agreement to what our indirect Seahorse measurements suggested.

### MitoTEMPO reverts CSE-driven PA vasodilation decrease, but does not recover normal vasoconstriction responses

After showing that mitochondrial superoxide chelator mitoTEMPO could indeed prevent some of the mitochondrial dysfunctions triggered by CSE in hPASMC *in vitro*, we then addressed the repercussions this could have on the PA functionality. If these alterations affecting the mitochondria network in hPASMC were mainly responsible for the lack of contractile force that CSE exerted on the PA, it could be reasonable to expect that mitoTEMPO might alleviate part of this stiffened phenotype. Following *ex vivo* exposure of murine PA, we observed that CSE blunted vasoconstriction responses to KCl and 5-HT and to a similar extent both in the absence and presence of mitoTEMPO (Figure 3A). However, mitoTEMPO did show beneficial effects upon recovering the CSE-driven loss of PA vasodilation capacity (Figure 3B). Although we only detected a slight, not significant recovery of endothelium-dependent vasodilation, elicited by ACh (Figure 3B, left), the effects on endothelium-independent vasodilation, triggered by SNP treatment, did reach statistical significance (Figure 3B, right). Thus, the reduction of mitochondrial superoxide levels by mitoTEMPO could help preventing, at the tunica media level, some of the deleterious effects of CSE on the vasodilation response of the PA.

**Figure 3:**
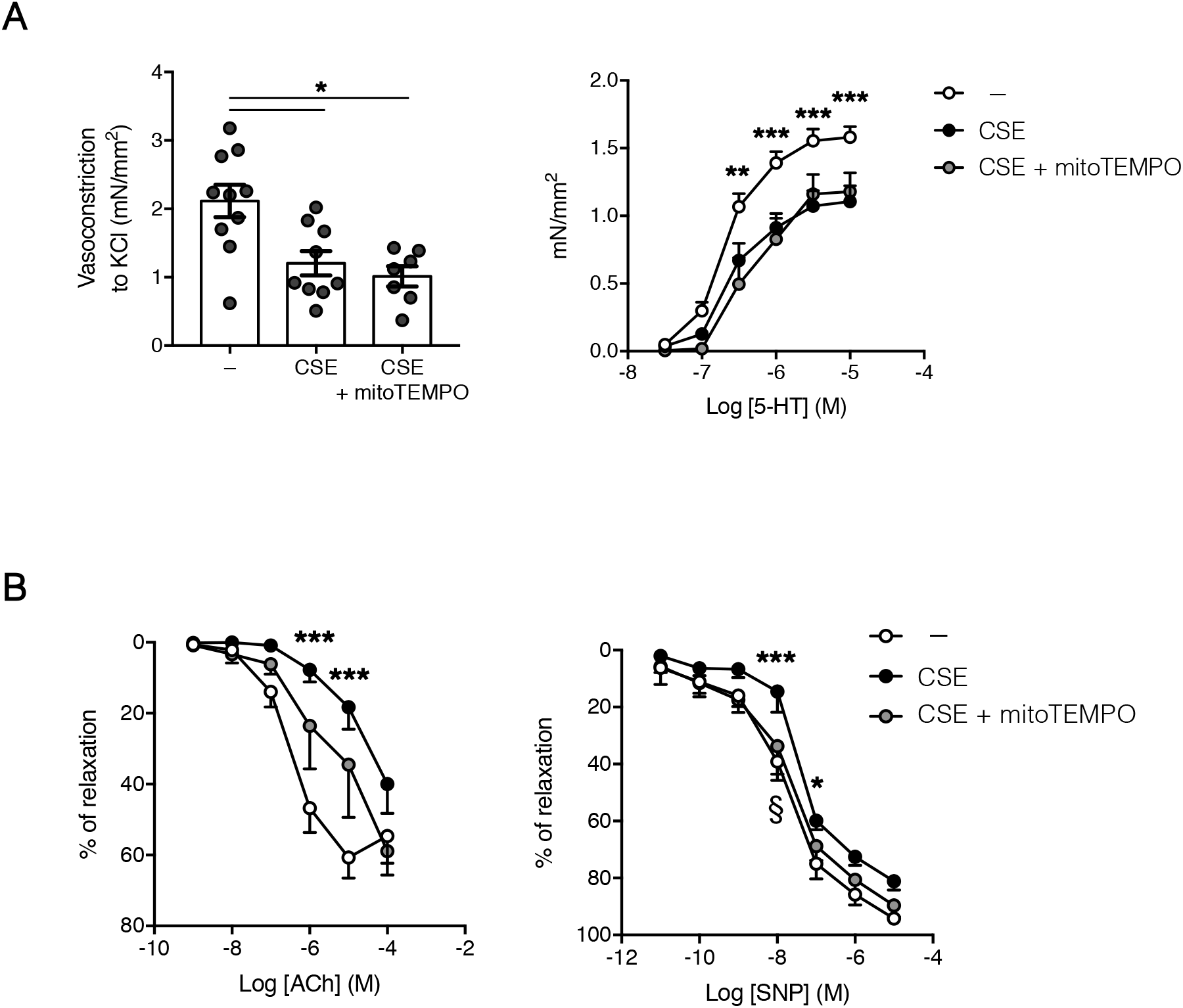
Vasoconstriction and vasodilation responses of PA challenged to CSE and mitoTEMPO. Following overnight *ex vivo* challenge with 10% CSE in the absence or presence of mitoTEMPO, vascular responses of endothelium-intact PA from WT C57BL/6J mice were analyzed by wire myography. **A.** Vasoconstriction responses to 80 mM KCl (left) or to increasing amounts of 5-HT (right) were presented as mean ± SEM; n = 7-10. Statistical comparisons among groups were made using Kruskal-Wallis’ *H* test followed by Dunn’s *post hoc* test (left, \**P*<0.05) or repeated measures two-way ANOVA test followed by Bonferroni’s *post hoc* test (right, \*\**P*<0.01, \*\*\**P*<0.005 between non CSE-exposed and CSE-exposed groups in the absence of mitoTEMPO). **B.** Vasodilation responses to increasing amounts of acetylcholine (ACh, left) or sodium nitroprusside (SNP, right) were expressed as percentages of relaxation related to 10 μM 5-HT-driven vasoconstriction, and presented as mean ± SEM; n = 4-7. Statistical comparisons among groups were made using repeated measures two-way ANOVA test followed by Bonferroni’s *post hoc* test (\**P*<0.05, \*\*\**P*<0.005 between non CSE-exposed and CSE-exposed groups in the absence of mitoTEMPO; ^§^*P*<0.05 between CSE-exposed groups in the absence and presence of mitoTEMPO).

### CSE treatment results in the downregulation of Cyb5R3 levels in hPASMC

Taking into account these previous results, we wondered if CSE-driven upregulation of mitochondrial superoxide could somehow interfere with the NO signaling through the sGC pathway, given that mitoTEMPO was able to restore PASMC sensitivity to the NO-donor SNP. We observed that CSE did not produce a significant decrease of the levels of sGC subunits (Figure 4A). It has been recently proven that sGC sensitivity to NO is regulated by Cyb5R3, which maintains the fundamental reduced status of the heme group in sGC, this allowing for proper NO signaling and downstream cGMP synthesis^61^. Our data showed that CSE, rather than affecting structural sGC subunits, produced a significant downregulation of Cyb5R3 protein levels (Figure 4B). This could imply that, in a superoxide-enriched scenario such as the one resulting from CSE treatment, antioxidant systems in charge of keeping sGC redox status might be insufficient, and sGC may become less sensitive to endothelial NO as well as to NO-donors like SNP.

**Figure 4:**
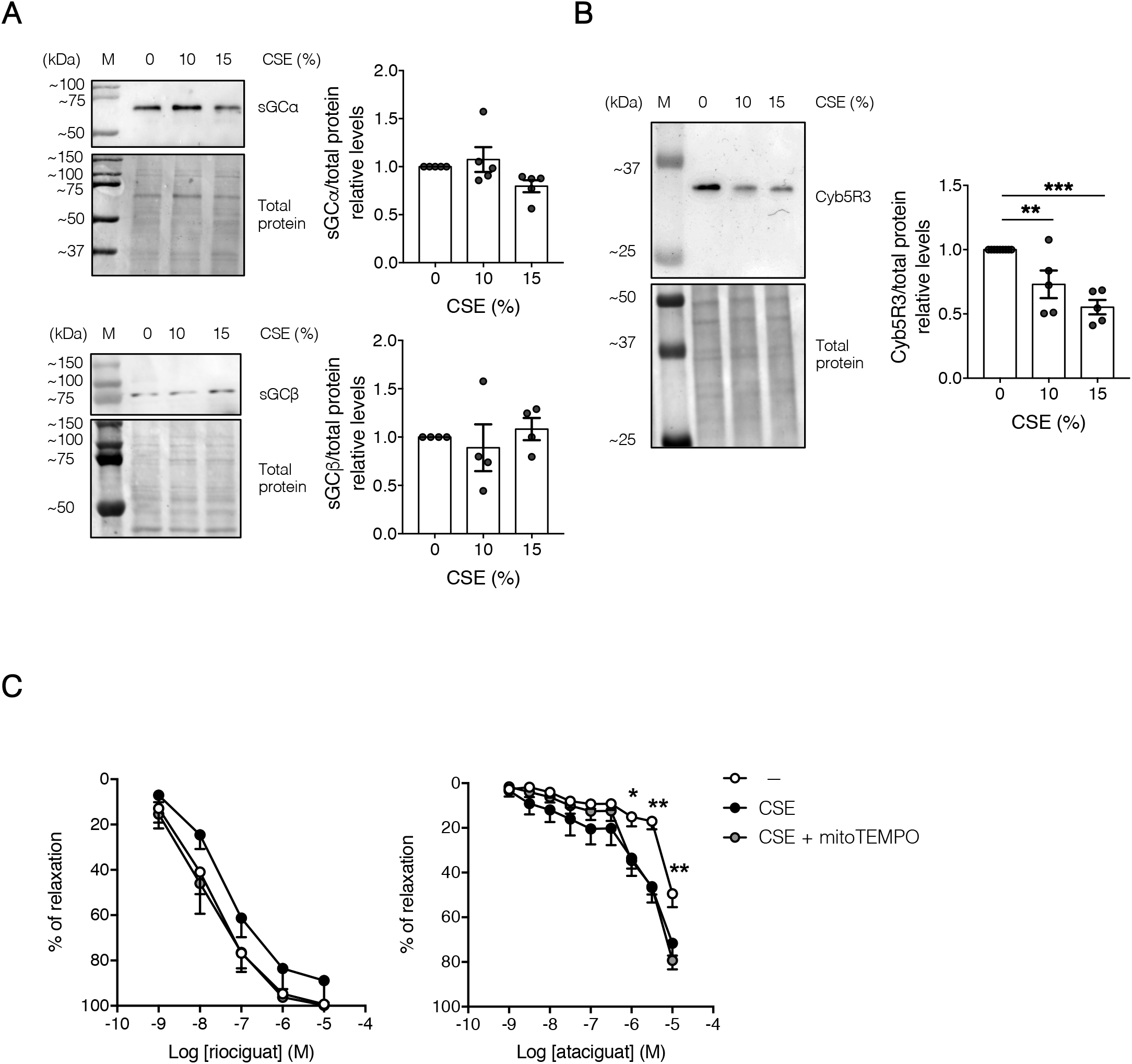
CSE impact on sGC subunits and Cyb5R3 enzyme levels in hPASMC. Following 24-hour challenge with CSE at the indicated concentrations, the expression of sGC subunits (**A**) and reductase Cyb5R3 (**B**) in hPASMC was analyzed by western blot. Protein levels of hPASMC were analyzed by western blot and probed against sGCα, sGCβ, Cyb5R3 and total protein. Representative images and band quantifications by densitometry are shown. M: molecular weight marker lane. Densitometry data were controlled with total protein staining as loading control, expressed as fold change over 0% CSE-exposed cells, and presented as mean ± SEM; n = 4-5. Statistical comparisons among groups were made using Kruskal-Wallis’ *H* test followed by Dunn’s *post hoc* test (Cyb5R3) or one-way ANOVA test followed by Bonferroni’s *post hoc* test (sGCα and sGCβ) (\*\**P*<0.01, \*\*\**P*<0.005). **C.** Following overnight *ex vivo* challenge with 10% CSE in the absence or presence of mitoTEMPO, vascular responses of endothelium-intact PA from WT C57BL/6J mice were analyzed by wire myography. Vasodilation responses to increasing amounts of the sGC stimulator riociguat (left) or the sGC activator ataciguat (right) were expressed as percentages of relaxation related to 10 μM 5-HT-driven vasoconstriction, and presented as mean ± SEM; n = 5-7. Statistical comparisons among groups were made using repeated measures two-way ANOVA test followed by Bonferroni’s *post hoc* test (\**P*<0.05, \*\**P*<0.01 between non CSE-exposed and CSE-exposed groups in the absence of mitoTEMPO).

### CSE-mediated increase on mitochondrial superoxide promotes sGC oxidation and diminishes the vasodilation response to riociguat

Oxidative stress within PASMC is closely related to the fundamental requirement of sGC heme group reduced status. Thus, we aimed to further explore whether this CSE-mediated decrease on Cyb5R3 was associated to an imbalance in the redox status of the sGC, in favor of a more oxidized form. To this aim we evaluated the PA vasodilation responses to the sGC stimulator riociguat, and sGC activator ataciguat. Our results indicated that sGC excessive oxidation might indeed be occurring, since we observed a CSE-mediated reduction of PA vasodilation response to riociguat while the response to ataciguat was instead increased significantly (Figure 4C). Interestingly, mitoTEMPO alleviated the CSE-driven reduced response to riociguat, whereas no clear changes in the response to ataciguat were observed (Figure 4C).

## DISCUSSION

COPD is a major health concern worldwide, and although the current therapies alleviate its symptoms, the disease is not definitely resolved. Therefore, it is necessary to unravel the mechanisms involved in COPD and COPD-related PH in order to design novel and effective treatments. Considering that oxidative stress is a major pathological trait of COPD^15–20^, we tried to better decipher its contribution to the alterations we previously observed on PA and PA isolated cells^8^. In this respect, it has been widely considered that alleviating oxidative stress could have a positive impact on the management of COPD patients, and therefore a revision of antioxidant treatments might enhance the benefits of currently approved vasodilation-targeted therapies. From a pharmacological perspective, this could widen the repertoire of molecular targets to address in COPD-related PH.

Our present results showed an increase in mitochondrial superoxide levels in hPASMC challenged to CSE. This increase was not counterbalanced by an increase in the endogenous antioxidant defenses, since these were unaltered or, even, downregulated. This could be explained in terms of *NFE2L2* mRNA downregulation, as well as insufficient NRF2 stabilization and nuclear translocation upon CSE treatment, as shown in COPD patients ^26,46-48^. Nevertheless, our data pointed towards NRF2 being indeed stabilized in hPASMC challenged to CSE. Thus, we cannot discard that additional defects on its transcriptional activity could explain why the highest dose of CSE did not result on increased antioxidant enzymes levels. Indeed, similar findings are reported to occur in bone marrow derived macrophages when CSE-exposed *in vitro*^76^. In these studies, Lee *et al* demonstrate that despite CSE stabilizes NRF2, this is accompanied by a downregulation of acetyltransferase p300, essential for its transcriptional activity^77^. Should this be the case in our PA cells, a high CSE dose-triggered p300 decrease might dampen NRF2 stabilization effects and impede a proper antioxidant defense upregulation.

Besides mitochondrial ROS levels, our experiments addressing their network structure after CSE challenge suggested a mitochondrial dysfunction, with a more fragmented network than control hPASMC. Accordingly, Aravamudan *et al*^74^ also found a CSE-driven profound disruption of mitochondrial network in human airway SMC with increased fission-promoting dynamin-related protein 1 levels. Most importantly, in agreement with our own results in PASMC, these fragmented mitochondria also have higher ROS levels and decreased respiration parameters, including ATP production. Similar hypoxia-driven mitochondrial alterations were recently shown on rat PASMC^78^. These authors demonstrate that hypoxia-driven fissioned mitochondria contribute to exert more vasoconstriction force and increased PAP, although, unfortunately, these mitochondria energetics were not addressed. Nevertheless, these data suggest that although hypoxia may induce a similar mitochondrial network fragmentation compared to CS, their repercussion onto PA contractility may not be identical.

Our recently published results demonstrate that high N-acetyl cysteine concentration, prevents CSE insults, particularly oxidative stress-triggered cell senescence induction^8^. Previous studies by Ito *et al* demonstrate that treating human bronchial epithelial cells with mitoTEMPO effectively diminishes mitochondrial ROS production triggered by CSE and reduces the upregulation of senescence markers^79^. Furthermore, other authors have shown that ozone-mediated pulmonary damage is also reverted by diminishing oxidative stress with mitoTEMPO in mice^71^. In our present study we evaluated the antioxidant effects of mitoTEMPO on PA dysfunction, and also observed a partial amelioration of CSE-driven damage.

Together with ROS burden and fragmented mitochondrial network, our data showing decreased OCR and depolarized Δψ_m_, might imply a CSE-driven deficient ATP production that might explain our recent published results (confirmed in this present study) in which CSE promotes the loss of contractile force at the PASMC level^8^. However, our present results only showed a modest decrease on total ATP levels due to CSE exposure, and mitoTEMPO did not prevent this. These results suggested that, because of the complex machinery needed for contractility generation, additional alterations affecting Ca^2+^-related signaling molecules, cytoskeleton proteins or ECM components might also contribute to this hypocontractile phenotype and therefore deserve a more systematic exploration in the future.

Similar to reduced vasoconstriction, we wondered if mitochondrial ROS would also contribute to exacerbate the vasodilation decrease we observe. Our data showing that mitoTEMPO recovered PA sensitivity to SNP support this idea, and could have a number of explanations worth exploring. On the one hand, excessive superoxide burden within PASMC likely targets NO-sGC vasodilator signaling pathway, because of the scavenging nature of superoxide on NO to form peroxynitrite^52^. This would not only diminish bioavailable NO for sGC activation, but could also contribute to exacerbate oxidative stress on PASMC. Indeed, our results showing a CSE-driven SOD3 downregulation might indicate poor extracellular antioxidant defenses in the PA cells vicinity, which could therefore contribute to NO scavenging. Besides remaining eNOS activity in PA tunica intima, inducible NO synthase (iNOS) should also be considered as a source of ONNO^-^. Gredic *et al* recently described that myeloid cell-specific iNOS-deficient mice are protected against chronic CS-driven pulmonary vascular muscularization, right ventricular systolic pressure and right ventricle hypertrophy, but still become affected by emphysema. These authors also found that iNOS deletion in macrophages abolishes their ability, upon CSE challenge *in vitro*, of stimulating PASMC proliferation and migration^80^. However, whether immune iNOS-derived NO contributes to oxidative stress in PA media is still to be evaluated.

On the other hand, oxidative stress within PASMC might be closely related to the fundamental requirement of sGC heme group reduced status. Our data suggest that the levels of the main enzyme maintaining this reduced form, Cyb5R3^61^, decreased following acute CSE challenge. In a pro-oxidant environment, this could translate onto an excessive sGC oxidation and, thus, diminished sensitivity to NO. This has actually been described in patients suffering from cerebral arteriopathy, whose vascular SMC indeed show significant sGC oxidation and reduced cGMP levels^81^. Our results indicated that this might in fact be the case, since we observed a better PA vasodilation response to the sGC stimulator riociguat in the presence of mitoTEMPO, this indicating that excessive mitochondrial superoxide burden within the PA prevented the vasodilation response due to an imbalance on the sGC redox state. These results are relevant and point out to the pharmacological benefits of vasodilation-related drugs in combination with specific sGC protection against the oxidative stress caused by CS, perhaps through mitochondrial ROS scavenging. In line with this, sGC activators might represent a preferred strategy compared to sGC stimulators under pathological conditions associated with enhanced sGC oxidation, such as chronic CS exposure.

In conclusion, our results demonstrate that CSE exposure causes an increase on superoxide levels in PASMC. We provide evidence of mitochondria as the most likely cellular source of increased ROS. In addition, this study shows an inadequate antioxidant response in CSE-treated PASMC, thus suggesting that oxidative stress is a crucial mechanism, which might have adverse effects in the pulmonary vasculature. Particularly, our data clearly show that CSE-challenged hPASMC display decreased levels of Cyb5R3 enzyme, which is involved in sGC redox homeostasis. Therefore, our results suggested that CSE-mediated increase of mitochondrial superoxide might contribute to compromise vascular responses in the PA. Given the increasing COPD burden nowadays, it is urgent to further characterize these pathways and consider mitochondrial oxidative stress alleviation in combination with stimulators of the NO-signaling pathway as a promising therapy for COPD-related PH recovery.

## ABBREVIATIONS

5-HT: serotonin
ACh: acetylcholine
ANOVA: analysis of variance
COPD: chronic obstructive pulmonary disease
CS: cigarette smoke
CSE: cigarette smoke extract
Cyb5R3: cytochrome b5 reductase 3
eNOS: endothelial nitric oxide synthase
FCCP: carbonyl cyanide-*p*-trifluoromethoxyphenylhydrazone
hPASMC: human pulmonary artery smooth muscle cells
iNOS: inducible nitric oxide synthase
NO: nitric oxide
NOX: NADPH oxidase
NOXA1: NOX activator 1
NOXO1: NOX organizer 1
NRF2: nuclear erythroid-2 related factor 2
OCR: oxygen consumption rate
PA: pulmonary artery
PAP: pulmonary artery pressure
PASMC: pulmonary artery smooth muscle cells
PH: pulmonary hypertension
ROS: reactive oxygen species
RS: recording solution
SEM: standard error of the mean
sGC: soluble guanylyl cyclase
SMC: smooth muscle cell
SNP: sodium nitroprusside
SOD: superoxide dismutase
TMRM: tetramethylrhodamine methyl ester
Δψ_m_: mitochondrial membrane potential

