## Supplemental figures and figure legends for "Cigarette smoke induces pulmonary arterial dysfunction through an imbalance in the guanylyl cyclase redox status"

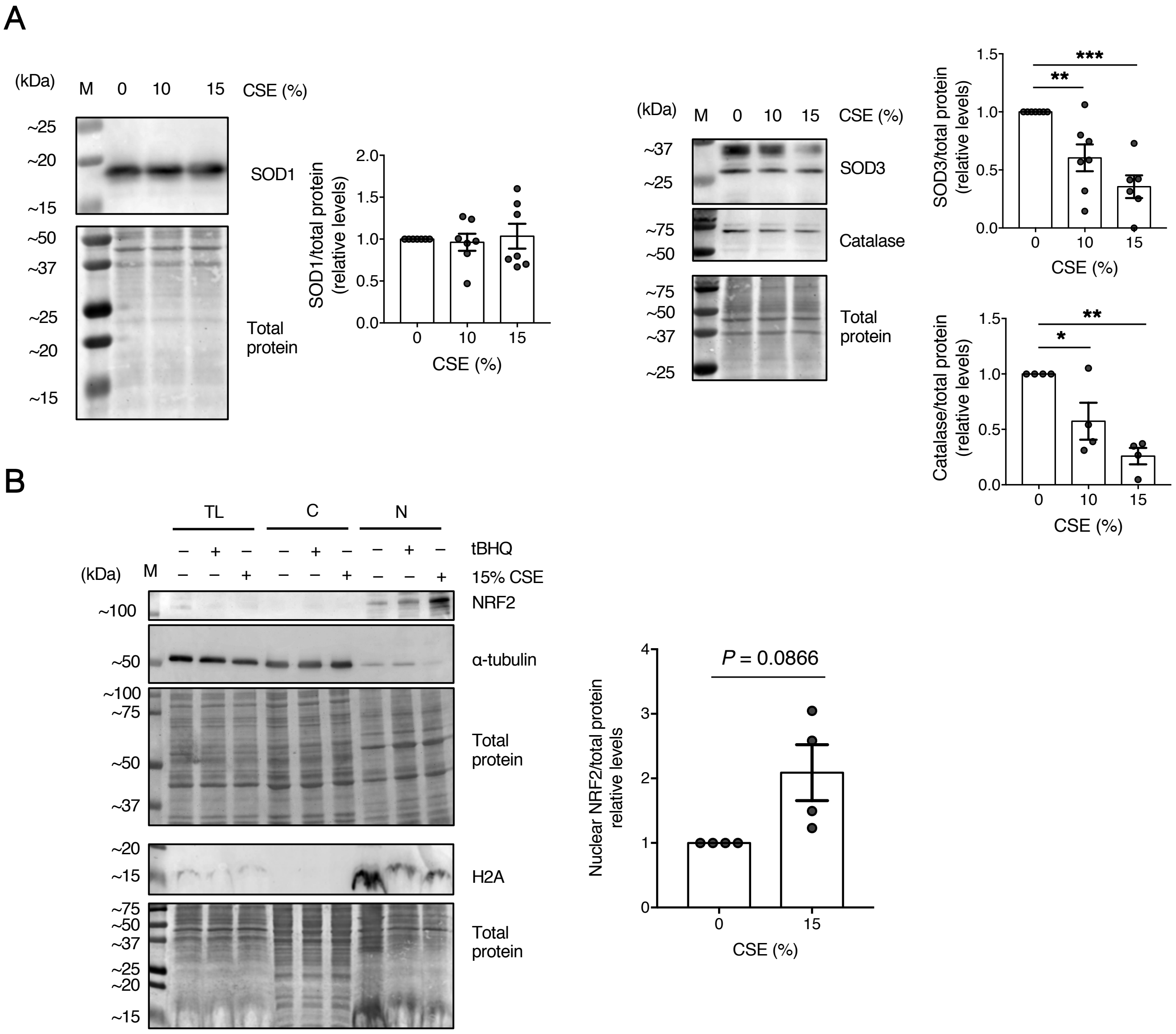


Supplementary figure 1

**Supplementary figure 1: Effect of CSE on endogenous antioxidant enzymes and NRF2 nuclear accumulation in hPASMC. A.** Following 24-hour challenge with CSE at the indicated concentrations, the expression of several antioxidant enzymes in hPASMC was analyzed by western blot, probed against SOD1, SOD3, catalase and total protein. Representative images and band quantifications by densitometry are shown. M: molecular weight marker lane. Densitometry data were controlled with total protein staining as loading control, expressed as fold change over 0% CSE-exposed cells, and presented as mean ± SEM; n = 5-7. Statistical comparisons among groups were made using one-way ANOVA test followed by Bonferroni's *post hoc* test (*$P$<0.05, **$P$<0.01, ***$P$<0.005). **B.** Following 24-hour challenge with 15% CSE or tert-butylhydroquinone (tBHQ) as NRF2 stabilization positive control, protein levels of total lysates (TL), cytosolic (C) and nuclear fractions (N) from hPASMC were analyzed by western blot. Cytosolic and nuclear extracts purity was assessed by probing the membranes against $\alpha$-tubuline and histone H2A. Representative images (left) and band quantifications by densitometry of nuclear NRF2 levels in control and CSE-treated hPASMC (right) are shown. M: molecular weight marker lane. Densitometry data were controlled with total protein staining as loading control, expressed as fold change over 0% CSE-exposed cells, and presented as mean ± SEM; n = 4. Statistical comparisons between groups were made using two-tailed one-sample Student's *t* test.

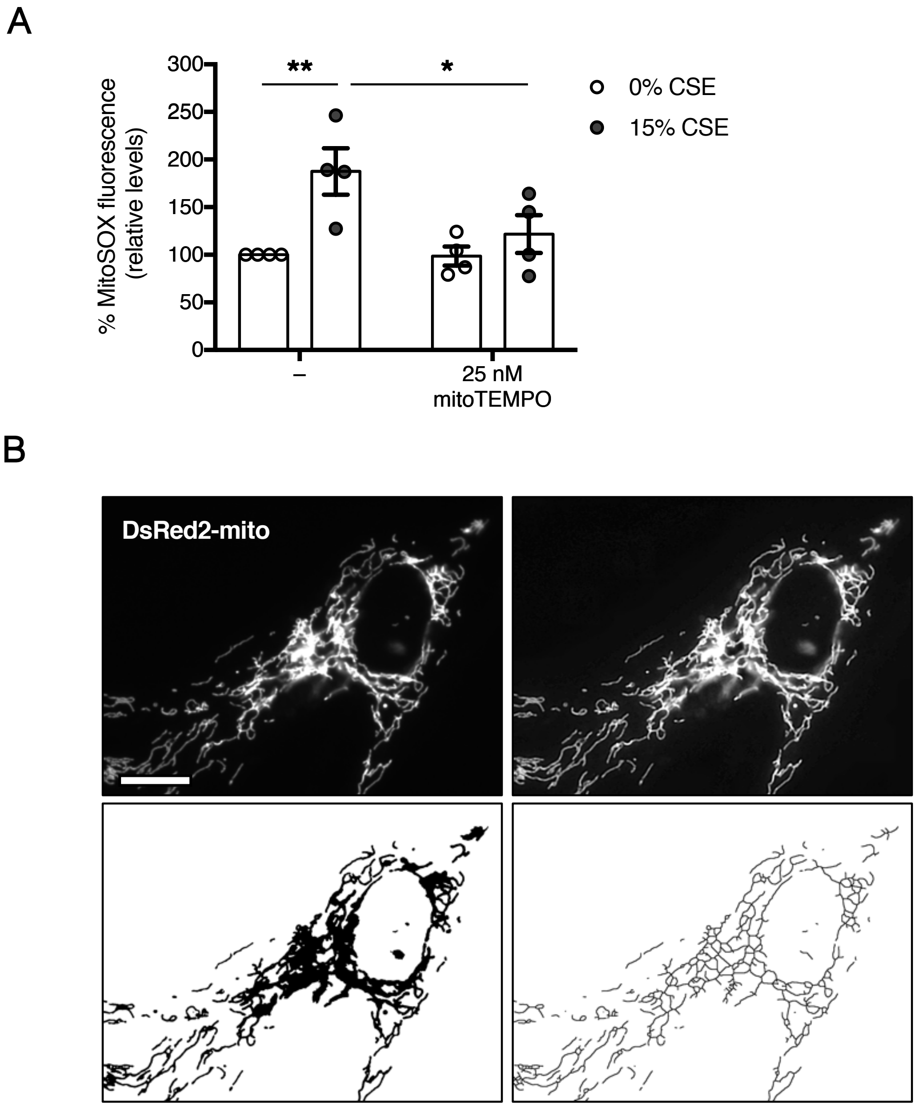


Supplementary figure 2

**Supplementary figure 2: Mitochondrial network image analysis. A.** Following 24-hour challenge with CSE in the absence or presence of 25 nM mitoTEMPO, mitochondrial $O_2^{\cdot -}$ levels in hPASMC were analyzed by flow cytometry, using 5 μM MitoSOX™. Values were calculated as the median fluorescence intensity of MitoSOX™-stained cells minus the median fluorescence intensity of unstained cells, expressed as percentages over 0% CSE-exposed cells, and presented as mean ± SEM; n = 4. Statistical comparisons among groups were made using two-way ANOVA test followed by Bonferroni's *post hoc* test (**P*<0.05, ***P*<0.01). **B.** Processing workflow of a representative image of pDsRed2-mito transfected hPASMC for mitochondrial network characterization. Representative raw image (top, left), processed image after unsharp masking, local contrast enhancing and median filtering (top, right), binary mask (bottom, left) and skeletonized representation of the mitochondrial network (bottom, right) are shown. Scale bar = 10 μm.
